# Differential Capture Modalities of Insect Traps Produce Contrasting Biodiversity Patterns

**DOI:** 10.1101/2025.09.17.676850

**Authors:** Thomas P. Franzem, Sophia Carmichael, Drew A. Davis, Katherine McNamara Manning, Christie A. Bahlai

## Abstract

Describing insect community and population trends is essential to their conservation. To assess insect populations and communities and identify potential drivers of change in insect populations, we often need to measure in a variety of ways and in a variety of ecosystems. Insect sampling is not standardized: data is frequently collected via different types of insect traps with different targets and modes of capture to sample the insect community. However, different traps may identify contrasting ecological signals, depending on their structure, deployment, and their effective range, and further, these differences may interact with the habitats they are deployed in. Here, we aim to assess the consistency of biodiversity patterns produced from different trapping methods deployed in array across several habitats. We investigated three trapping methods: an ant bait trap, a malaise trap, and a fermentation trap. We deployed these traps in a suburban forested area and at a restored forest. We computed richness and relative abundance of insects captured in each trap and assessed community compositional differences between traps and habitats. We found that each trap type produced differential patterns of richness, abundance, and composition from each other, and these patterns also varied between sites for most trap types. The contrasting patterns observed among traps is likely due to differing effective capture ranges of each trap; traps with smaller effective capture ranges were more sensitive to local habitat conditions than traps with broader effective capture ranges.

## Introduction

Insects are ecologically and economically important organisms (Losey & Vaughan, 2006; Jankielsohn, 2018; Verma et al., 2023) and occupy diverse ecological roles as pollinators, decomposers, herbivores, predators, and more (Noriega et al., 2018; Crespo-Pérez et al., 2020). The past several decades has seen declines in insects across taxonomic and functional groups, which threaten to disrupt the valuable ecosystem services they provision (Hallmann et al., 2017; Wepprich et al., 2019; Wagner, 2020; Halsch et al., 2025). Recent trends of global insect decline are concerning and have potentially far-reaching impacts on human-society (Wagner, 2020; Hailay Gebremariam, 2024). Insect declines are driven by multiple purported drivers, however precise inference concerning how, why, and when different insect groups are declining remains elusive due to the high diversity of insects and differential responses to habitat conditions (Hailay Gebremariam, 2024; Halsch et al., 2025).

Investigating insect population patterns, across groups and in different habitat contexts is necessary to identify drivers of declines and inform conservation action (Grames et al., 2022; Cooke et al., 2025). However, collecting quality insect data from naturally occurring insect populations across environmental conditions is challenging (Van Klink et al., 2022). There are multiple interacting factors that produce the observed data, which can introduce biased data and the detection of spurious or inaccurate trends (Zhang et al., 2021; Díaz-Calafat et al., 2024). For example, the length of a given study can produce inaccurate results (Bahlai et al., 2021), or the trap type used can create bias in the taxa represented in the dataset (Busse et al., 2022). Indeed, choices made by the researcher interact with stochastic natural processes (MacKenzie et al., 2002), and sometimes even very subtle changes in methodology can lead to detectable differences (McNamara Manning et al., 2024). The result of this interaction is that resultant data may not fully capture the ecological state being investigated or may mask changes in dynamics. In the context of insects, different traps sample different components of the insect community, with varying levels of specificity, based on their interactions with insect behavior, habitat use and movement. For example, malaise traps are a flight intercept trap that broadly sample any flying insects moving through a given area (Uhler et al., 2022). Conversely, pheromone lures specifically attract particular insects responding to often species-specific semiochemical cues (Fountain et al., 2014; Hanks et al., 2016). Thus, the observations drawn from biodiversity or community ecology studies are likely to be methods sensitive – that is, the ultimate patterns extracted from the data are shaped by the methods researchers use (McNamara Manning et al. 2022). Because different traps capture different insects, different experimental contexts and research questions will employ different collection methods, but the question remains - how sensitive is the overall biodiversity or community *pattern* to the method? For example, a study investigating bee community ecology used both sweep netting and bee bowls to sample the bee community (Pei et al., 2022). The authors identified contrasting patterns of richness, composition, and abundance from the different sampling methods (Pei et al., 2022). Similarly, an assessment of the efficacy of different common monitoring methods for crop pests found substantial differences in the abundance and richness of insects among methods (Gill & O’Neal 2015). Identifying how data produced from insect traps with different levels of taxonomic specificity and capture modalities influences the conclusions made about the system is a salient area of research given the need for more insect data to assess and study insect declines. Indeed, the results of insect biodiversity studies will be used to extrapolate to broader trends in insect populations and inform conservation and management (Saunders et al., 2019; Vallecillo et al., 2021), so more context is necessary to identify how trap modality impacts the observed patterns of insect communities and populations.

Here we aim to assess the biodiversity patterns produced by different insect traps deployed in habitats with similar structure but differing landscape context. Specifically, we address the question: do insect traps that are functionally different in their method of capture identify the same patterns across sampling sites? To address this question, we employed three insect trap types that differ in capture mode and specificity. We deployed these traps at two site complexes that differed in current land use and compared the communities captured from each trap type at each site complex. Results of this work can provide insight into experimental design and optimize insect ecology surveys.

## Methods

### Study Area

We collected insects at two site-complexes in Northeast Ohio, U.S.A., with each site complex containing two sampling locations. One site-complex was on Kent State University’s campus, and the other was at the Snowville Restoration Site in Cuyahoga Valley National Park (Fig 1; National Parks Service collection permit CUVA-2023-SCI-0011). We selected these two locations because they differ in their contemporary land use and surrounding landcover and could be sampled regularly over the course of the summer season, to align with the availability of labor. The KSU campus is a landscaped, suburban ecosystem, whereas the Snowville Restoration Site is a mix of forest and open habitat and is surrounded mostly by natural and regenerated landcover in Cuyahoga Valley National Park. Each sampling location within a site-complex was a minimum of 500 m from the other sampling location. At each sampling location, we haphazardly selected a point for the malaise trap, fermenting bait trap, and three ant-bait traps (see below for trap details). Ant-bait traps were at least three meters away from other traps, and malaise and fermentation traps were set a minimum of 10m apart. Sampling occurred weekly from June 24th through July 26^th^, 2024. Each trap type had different exposure times due to the different modalities of the traps, thus the number of samples among trap types varied (see below for details).

**Figure 1:**
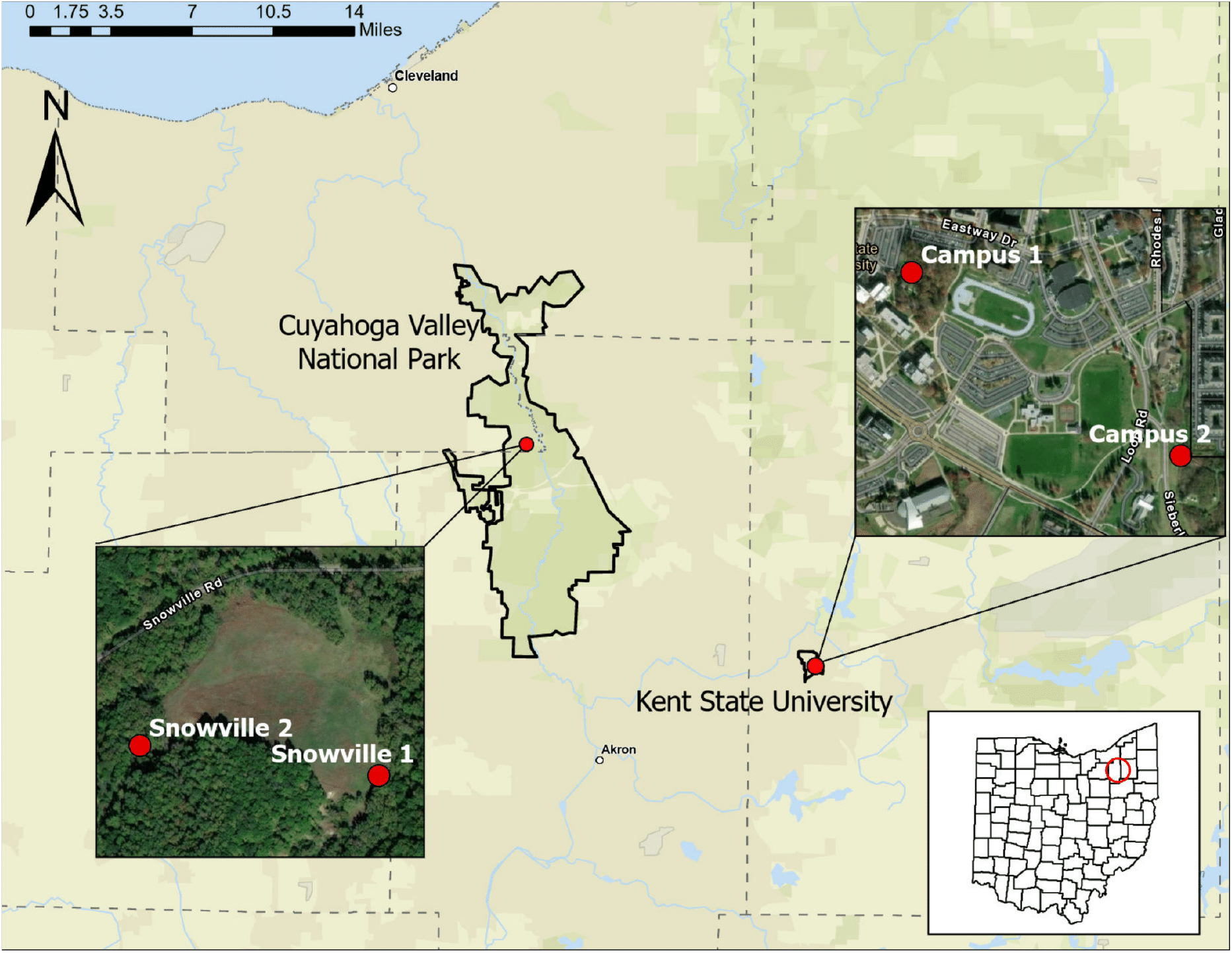
Site complex locations within northeast Ohio, USA (inset map). Each site complex contained two sampling locations. The ‘Snowville’ site complex was located in Cuyahoga Valley National Park, and the ‘Campus’ site complex was located on Kent State University’s main campus.

### Insect Traps

We used malaise traps, fermenting bait traps, and ant-bait traps because these traps range in capture mode, effective range, and specificity. Malaise traps capture any insects that fly, fermenting bait traps target insects that respond to specific olfactory cues, and ant-bait traps specifically attract ants. Further, fermenting bait traps have a relatively large effective range due to their longer exposure time and constant emission of attractive chemical plumes, malaise traps have smaller effective ranges because they are a passive flight intercept trap and thus only collect insects that are occupying and moving through the area, and ant bait traps have a small effective range because of their short exposure time and the relatively weak signal from the bait.

#### Malaise Traps

We used Townes Style Malaise Traps (Model BT1011) from MegaView Science Co., Ltd and utilized a wet-collection head with propylene glycol as the preservative. Malaise traps were set for three days per site, every other week, for a total of three malaise samples across the study period. Once specimens were returned to the lab, we transferred them to 70% ethanol for long-term storage.

#### Fermenting Bait Traps

Fermenting Bait Traps followed the bait recipe detailed in MacRae and Rice (2007) and MacRae (2015). We specifically used the beer-based formulation, and did not employ the wine-based fermenting bait detailed in MacRae (2015) due to logistical constraints. In brief, the bait was a diluted molasses mixture combined with 4.5g of active yeast and one 12 oz beer. Bait was mixed in a one-gallon bucket in the field, and the bucket was hung approximately 2 meters off the ground from a tree. Trap contents were collected weekly for a total of four sampling periods. Trap contents were collected by pouring the bait through a fine-mesh sieve. We then transferred the sieved insects into an enamel pan and used forceps to remove debris and dirt. In the field, we doused specimens with DI water to remove excess fermentation fluid on the specimens and then transferred the insects into 70% ethanol. The bait mixture was replaced every 14 days, as per MacRae (2015).

#### Ant Bait Traps

Ant bait traps consisted of half of a Keebler’s Pecan Sandies Shortbread cookie (Kaspari, O’Donnell & Kercher, 2000; Ohyama, King & Jenkins, 2020; Roeder et al., 2021) in an uncapped 50 mL plastic vial that was placed on the ground for one hour to allow time for ants to discover and recruit workers to the bait. Each site had three ant bait traps that were set out five times over the course of our study period. We collected ant bait traps by capping the vials. We returned vials to the lab and freeze-killed specimens by placing the vials in a freezer. After 48 hours in the freezer, we transferred specimens to 70% ethanol for long-term storage.

### Sample Processing

Once returned to the lab, we washed all malaise and fermentation trap samples with DI water to remove remaining collection fluids. Bycatch (such as spiders or slugs) were recorded but specimens were not retained. The same level of taxonomic identification was not possible across trap types due to the different capture modalities of each trap. For example, malaise and fermentation traps produced a high volume of specimens across multiple orders, whereas the ant-bait traps produced relatively few specimens within a single family (Hymenoptera: Formicidae). As such, we processed specimens to the appropriate taxonomic resolution for each trap to address our study question. Specifically, malaise and fermentation trap samples were identified to order, while the ant-bait samples were identified to the lowest taxonomic resolution possible.

### Data Analysis

#### Richness and relative abundance

We enumerated the number of specimens of each insect order from the malaise and fermentation traps and the number of specimens of each ant species from the ant-bait traps. At each site we calculated order richness from the malaise and fermentation traps and ant species richness from the ant bait trap with the *specnum* function in ‘vegan’ (Oksanen et al 2024). For downstream analysis, we analyzed only the five most observed insect orders from the malaise and fermentation traps (Lepidoptera, Coleoptera, Diptera, Hymenoptera, and Hemiptera). Several other orders were detected in these traps but were only comprised of one to two individuals (i.e. Blattodea, Mantodea). We calculated relative abundance for the five most observed orders collected in the malaise and fermenting bait traps and the relative abundance of all the ant species collected. Further, we calculated the relative abundance of insects detected in each trap type, at each site. To assess the differences in the abundance of each order captured in each trap type, we conducted a Welch two-sample t-test with log-transformed abundance data to meet assumptions of normality. Similarly, we conducted a Welch two-sample t-test to assess differences in the abundance of each order captured at each site complex using pooled data from both traps.

#### Community composition among traps and site complexes

To assess the composition of the insect community detected using each collection method and among sampling locations, we subjected the dataset to Nonmetric Multidimensional Scaling (NMDS). We used the ‘metaMDS’ function from the R package ‘vegan’ (Oksanen et al 2024). We subjected different subsets of the data to NMDS to investigate patterns that emerge when using one trap type or the other. We generated NMDS plots using just the malaise trap data and just the fermenting-bait trap data, each with sampling location as the grouping variable. With the ant data, we separately constructed an NMDS with sampling location as the grouping variable.

## Results

### Richness and relative abundance

At each sampling location, the orders Coleoptera, Diptera, Hemiptera, Hymenoptera, and Lepidoptera were the most frequently detected. Total insect order richness was highest at Snowville 2 (Table 1) due to the detection of Blattodea, Mantodea, Orthoptera, and Psocodea, however these orders were singletons and were excluded from downstream analysis. Fermentation traps accounted for approximately half of the insects detected at each site, followed by malaise traps, and then ant-bait traps constituted a small percentage of total insects captured per site (Table 1). In both the malaise and fermentation traps, Diptera was the most observed order at each sampling location (Table 1). The fermenting bait trap primarily caught Coleoptera and Diptera while the other orders had low relative abundance detected at each sampling location (Table 1). Conversely, the malaise trap captured relatively few beetles, but caught more specimens from Hemiptera, Hymenoptera, and Lepidoptera than the fermentation trap (Table 1).

**Table 1:**
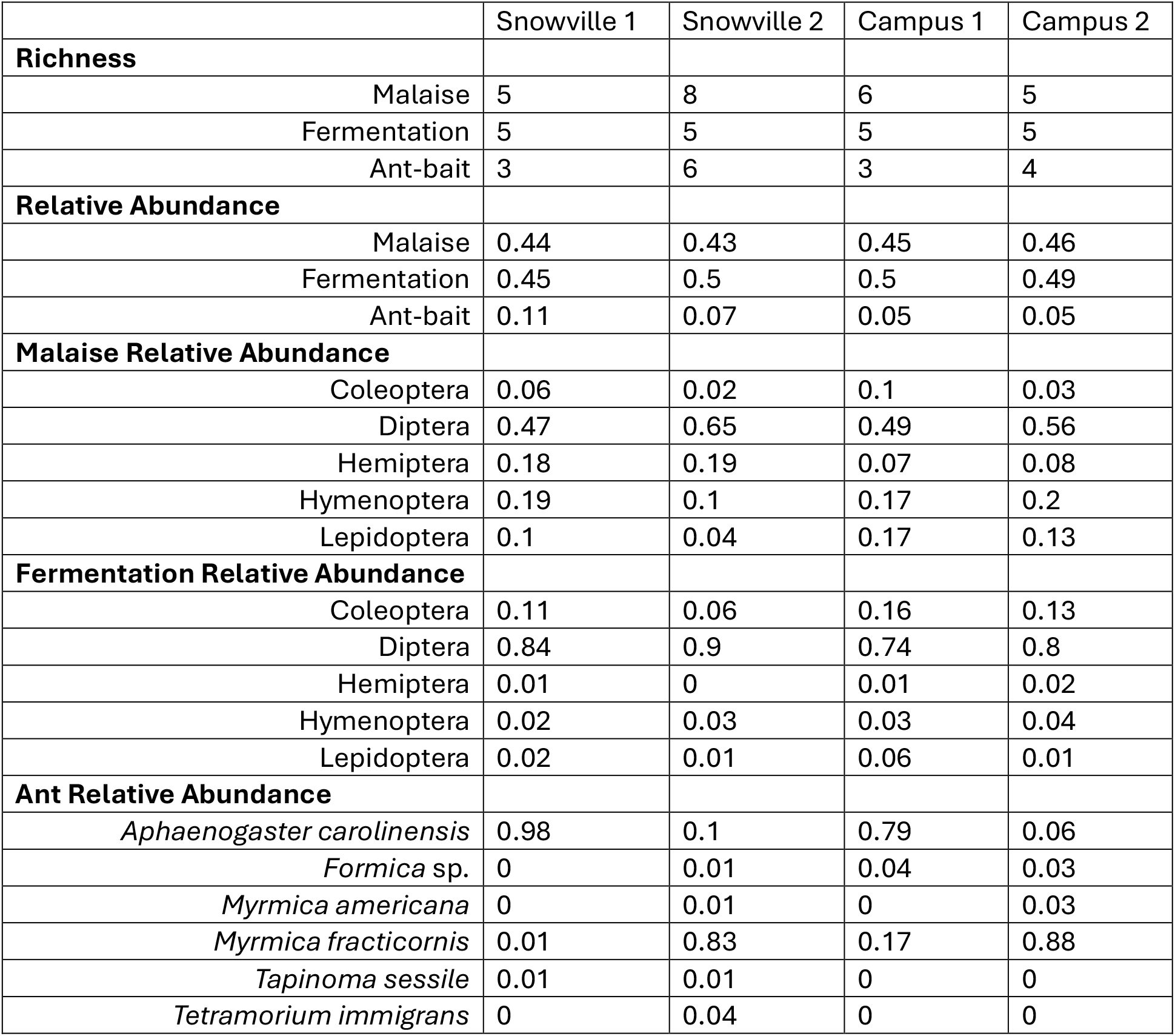
Richness of orders (Malaise and Fermentation) and species (Ant-bait) at each sampling site, as well as the relative abundance of both the total specimens collected and each order/species detected by each trap type at each site.

For ant species richness, we detected the highest ant richness at the Snowville 2 sampling location (Table 1). *Tetramorium immigrans* was only detected at Snowville 2 and *Tapinoma sessile* was only detected at the Snowville site complex. At Campus 1 and Snowville 1, *Aphaenogaster carolinensis* was the most observed species, and at Campus 2 and Snowville 2 *Myrmica fracticornis* was the most observed species (Table 1). All other ant taxa were detected at low abundances at each site (Table 1).

### T-tests between trap types and site complex

For each of the five most observed orders, we found a significant difference in the log abundance of orders detected between malaise and fermentation traps (Table 2). Specifically, the fermentation trap collected significantly more beetles and flies than the malaise trap, while the malaise trap collected significantly more true bugs, wasps and bees, and butterflies than the fermentation trap. When we compared the differences in order based on site complex, we found that for the fermentation traps there was no significant difference in the log abundance of each order between the two site complexes (Table 2). The malaise trap detected a significant difference in log abundance between site complexes only for true bugs (Table 2), and both traps combined did not detect a significant difference between site complexes for each order (Table 2).

**Table 2:** Results of t-tests performed for each order. We conducted t-tests to test for differences in order log-abundance (N) between trap types and site complexes.

|  | **Coleoptera** | **Diptera** | **Hemiptera** | **Hymenoptera** | **Lepidoptera** |
| --- | --- | --- | --- | --- | --- |
| **Fermentation** |  |  |  |  |  |
| Mean Log N | 3.66 | 5.7 | 1.14 | 2.38 | 1.84 |
| **Malaise** |  |  |  |  |  |
| Mean Log N | 1.97 | 5.23 | 3.62 | 4.02 | 3.35 |
| **T-test p-value** | <0.0001 | 0.018 | <0.0001 | <0.0001 | 0.003 |
| **Fermentation** |  |  |  |  |  |
| Campus Mean Log N | 3.72 | 5.48 | 1.47 | 2.36 | 2.11 |
| Snowville Mean Log N | 3.6 | 5.92 | 0.81 | 2.4 | 1.61 |
| **T-test p-value** | 0.618 | 0.114 | 0.063 | 0.884 | 0.55 |
| **Malaise** |  |  |  |  |  |
| Campus Mean Log N | 2.39 | 5.06 | 2.94 | 3.98 | 3.51 |
| Snowville Mean Log N | 1.65 | 5.4 | 4.29 | 4.06 | 3.18 |
| **T-test p-value** | 0.292 | 0.217 | 0.004 | 0.753 | 0.488 |
| **Both Traps** |  |  |  |  |  |
| Campus Mean Log N | 3.1 | 5.3 | 2.1 | 3.05 | 2.98 |
| Snowville Mean Log N | 2.57 | 5.7 | 2.3 | 3.11 | 2.4 |
| **T-test p-value** | 0.253 | 0.055 | 0.726 | 0.874 | 0.379 |

### Community composition

When we examined just the malaise trap data with ‘collection site’ as the grouping variable, we found the ‘campus’ sites were distinct from the CVNP sites (Fig 2a), however, when we examined just fermentation trap data, we found no difference in the communities detected (Fig 2b). When examining the ant communities, we detected different patterns relative to the malaise and fermentation trap data (Fig 2c). We found that collection sites within site complexes were separate, with one of the campus sites overlapping with both CVNP sites. We note, however, that few ants were detected at both campus sites, which may cause artefacts in NMDS construction. We conducted an NMDS using ‘site complex’ as the grouping variable and found similar patterns for each collection method as we did when using ‘sampling location’ as the grouping variable (Appendix: FigS1).

**Figure 2:**
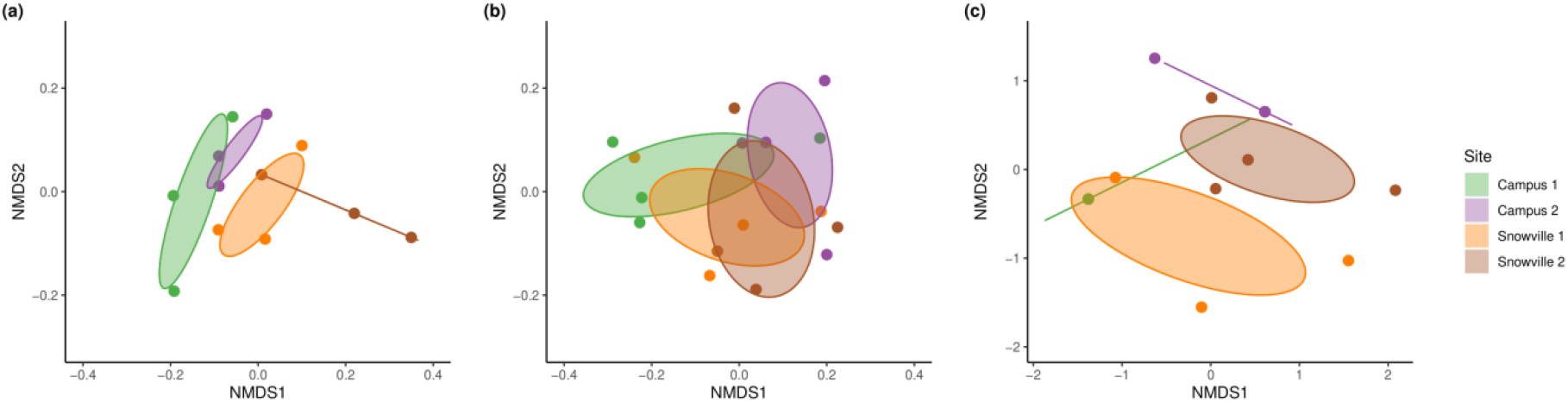
Non-metric multidimensional scaling representing insect community composition at four collection sites, captured in three different trap types. (a) Malaise trap: stress = 0.07, (b) Fermentation trap: stress = 0.11, (c) Ant bait trap: stress = 0.03. Ellipsoids represent 95% confidence interval of the mean for collection sites. Points displayed represent community composition for each sample.

We constructed NMDS plots with different grouping variables. With ‘trap type’ as the grouping variable, we found that the community of insects detected by the malaise traps and fermentation traps were different, based on the non-overlap of the standard deviation ellipses (Appendix: Fig S2). When we set ‘collection site’ as the grouping variable for the combined malaise and fermentation trap data, we found no difference in the communities detected among sites, as indicated by the overlap of standard deviation ellipses (Appendix: Fig S3).

## Discussion

We identified contrasting patterns of richness, abundance, and composition at each site complex, and found that these patterns varied with trap type. Although the sample size of this dataset is limited, our results demonstrate that different conclusions about the insect community at each site could be reached depending on the trap used. The similar proportion of insects captured from each trap type at each sampling location increases our confidence in these results, because this indicates the traps were performing consistently among sampling locations, and thus differences detected in composition, richness, and abundance was due to actual differences in the insect community.

Our observations align with the findings of Busse et al (2022), wherein they detected differential biodiversity patterns across trap types, but moreover these differences interacted with sampling location, creating differential patterns in biodiversity observations depending on sampling modalities. Our results suggest that the consideration of research context is essential to interpretation in insect biodiversity studies. Indeed, correcting for trap biases can facilitate more nuanced interpretation to insect ecology studies. For example, if researchers using our dataset were basing inference on data from just fermentation traps, they may conclude there is no difference in the insect community based on order richness (Table 1), abundance per order at each site complex (Table 2), and the composition of the community observed (Fig 2 b). Conversely, if instead the researchers were basing inference on just the malaise trap data, they may conclude the Snowville site complex is different than the KSU site complex, where order richness is higher at Snowville (Table 1), hemipterans are significantly more abundant (Table 2), and the communities are different based on the non-overlap of the standard deviation ellipses in the NMDS (Fig 2a).

Among traps we did not see patterns in richness, abundance, or composition converge as trap specificity increased. For example, when looking specifically at composition among sites, the pattern is different for each trap type; the most non-selective trap (malaise traps) found the site complexes were different based on NMDS, then the more specific fermentation trap found no difference, and the ant-bait trap NMDS plot identified patterns entirely different than the malaise or fermentation traps. A likely explanation for these contrasting patterns between trap types is the variable ‘effective range’ of the traps. The malaise and ant-bait traps have smaller effective ranges than the fermentation traps and are thus likely more sensitive to local-habitat conditions. Malaise traps do not use attractants, so capture is limited to insects already occupying and moving through the area. Similarly, the ant-bait traps are small and have a limited exposure time (1 hour), so only ants that are in the immediate vicinity will be attracted to the lure and have time to locate the bait and recruit workers. Conversely, the fermentation trap persistently emits a chemical plume that draws insects in from the surrounding area, thus differences in the insect community at the sampling locations may be overridden by the attractive force of the bait. With this result in mind, careful consideration should be used in planning insect surveys so that the scale of the study is matched by the scale of the trap.

Many insect biodiversity surveys are shifting towards making inference at coarser levels of taxonomic resolution (Kallimanis et al., 2012; De Oliveira et al., 2020) - that is order and family as opposed to species because of the desire to extend observations to general responses across related taxa. Focusing at this level of identification can facilitate more biodiversity research by decreasing the time, resources, and expertise needed to process samples (Obrist & Duelli, 2010; Timms et al., 2013; Carreira-Flores et al., 2024). Our results demonstrate that differences in the communities present among habitats can be detected when specimens are identified to the order level. This finding can inform the design of studies aiming to investigate how different habitats influence insect communities. An interesting avenue of future research would be to explicitly test how identifying samples to different levels of taxonomic resolution alter the patterns detected among habitats, and could provide an opportunity to examine functional traits, such as insect mobility (McNamara Manning et al., 2022).

In designing insect biodiversity studies, both the spatial and taxonomic scale at which researchers aim to make inference will inform the selection of traps. Our results demonstrate that the effective range of the traps can change the patterns detected, and thus careful consideration should be taken to ensure the spatial scale of the study matches the spatial scale of the trap. Malaise traps appear to offer a reasonable middle-ground for traps. They broadly sample the insect community occupying a site, but are sensitive enough to detect differences in community composition among sites. Furthermore, although they capture a significant volume of specimens, the number of specimens is proportionally lower than the number of specimens captured by the fermentation trap, so if researchers aimed to identify specimens to finer taxonomic resolution, it would be a manageable number of specimens. Conversely the fermentation traps, although selecting for a more specific component of the insect community, may have too large an effective range to detect fine-scale differences in community composition among sites, and thus are likely better suited for generating specimens for species inventories and investigating broader, regional trends. Given the sensitivity of ants to environmental conditions, and the relatively few specimens produced from the ant bait traps compared to the malaise and fermentation traps, ant bait traps are the most effective and efficient of the traps we tested here. There is growing focus among entomologists to reduce the ecological impact of biodiversity collections (Lövei et al., 2023; Barrett & Fischer, 2024); based on sheer abundance of specimens extracted, the ant-traps have the lowest ecological impact of the traps tested.

## Conclusions

We conclude that insect community patterns detected vary with trap type, likely due to differing effective capture ranges of each trap; traps with smaller effective capture ranges were more sensitive to local habitat conditions than traps with broader effective capture ranges. Future syntheses and metanalyses investigating insect biodiversity should account for the differential patterns that may emerge from the selected method. This is a salient line of inquiry that has not been extensively explored and should be more thoroughly investigated to properly contextualize insect biodiversity research (Busse et al 2022). Indeed, given the high diversity of insects, it is not surprising that traps targeting different components of the insect community and acting at different spatial scales produce different observed patterns. An interesting takeaway, though, is the apparent contrast in community composition we see in ants compared to the community composition of insect orders we detected from the malaise and fermentation trap. Ants are particularly sensitive to both biotic and abiotic conditions, which makes them a commonly used ‘ecological indicator’(Andersen & Majer, 2004; Lawes et al., 2017; Tiede et al., 2017). As stated above, because of the effective range of the ant traps, they are particularly sensitive to local habitat conditions. Thus, the patterns we detected among sites in the ant data may reflect fine scale and nuanced responses to habitat. We did not collect ants intensively enough or characterize site-specific environmental conditions at a fine enough resolution to specifically test what is driving the differences in the observed ant communities, however, our findings support the use of ants as fine-scale indicators of insect responses to environmental conditions.

## Acknowledgements

This research was conducted on Kent State University’s Kent Campus and in Cuyahoga Valley National Park (National Parks Service collection permit CUVA-2023-SCI-0011). Thanks to Leo Kuck for assistance with specimen collection, Robert Royd for assistance coordinating field work logistics, and the entire Bahlai Lab. This work was completed with support from Kent State University’s Summer Undergraduate Research Experience Program through the Office of Student Research and the National Science Foundation Faculty Early Career Development Program from the Division of Biological Infrastructure (DBI 2045721) to CB.

## Author Contributions

TF, CB, and KMM conceptualized the project. SC, DD, and TF performed data collection. SC performed initial specimen processing and identified ant species. DD enumerated the number of individuals detected from each insect order at each site. TF led the data analysis and figure creation with input from KMM, SC, and DD. SC generated figure 1. TF wrote the initial manuscript draft, based on posters made by SC and DD. All authors participated in manuscript revisions and editing.

