## Supplemental Figures 1, 2, and 3 for "Differential Capture Modalities of Insect Traps Produce Contrasting Biodiversity Patterns"

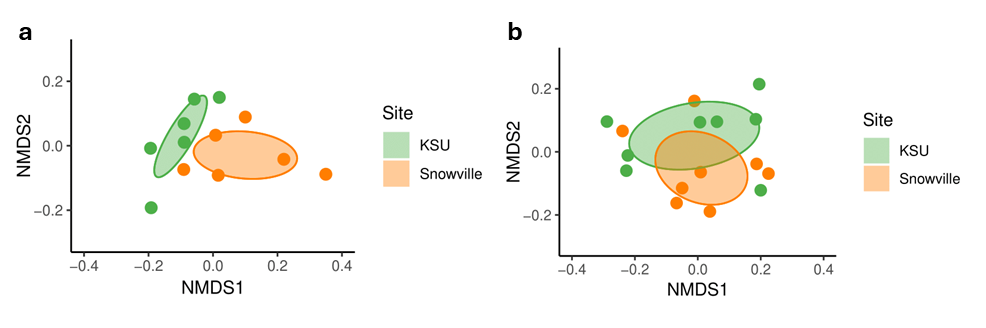

**Figure S1:** Non-metric multidimensional scaling representing insect community composition at site complexes, captured in (a) Malaise trap and (b) Fermentation trap. Ellipsoids represent 95% confidence of the mean for site complex. Points displayed represent community composition for each sample.

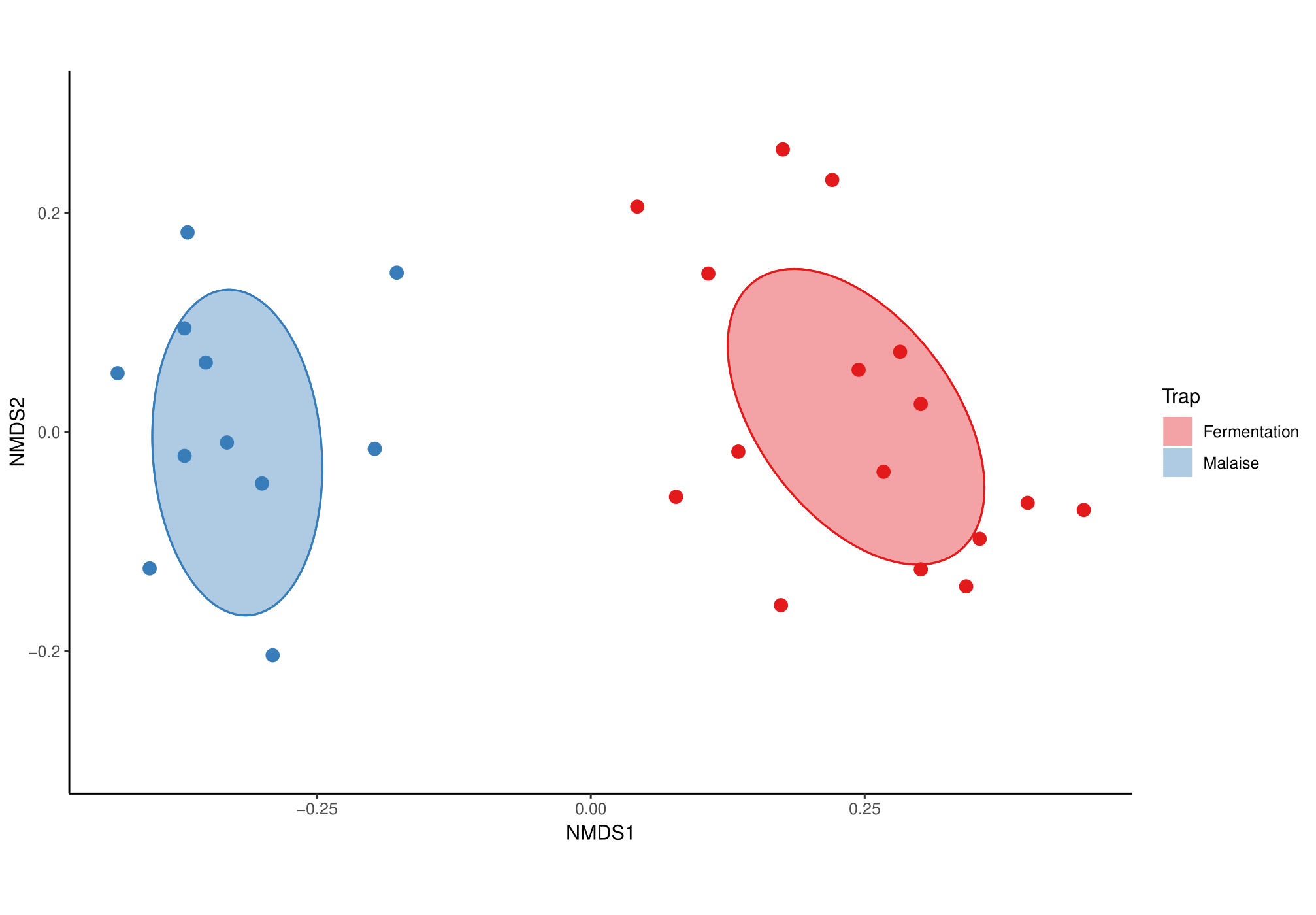

**Figure S2:** Non-metric multidimensional scaling representing insect community composition between fermentation and malaise traps. Ellipsoids represent 95% confidence of the mean for trap type. Points displayed represent community composition for each sample.

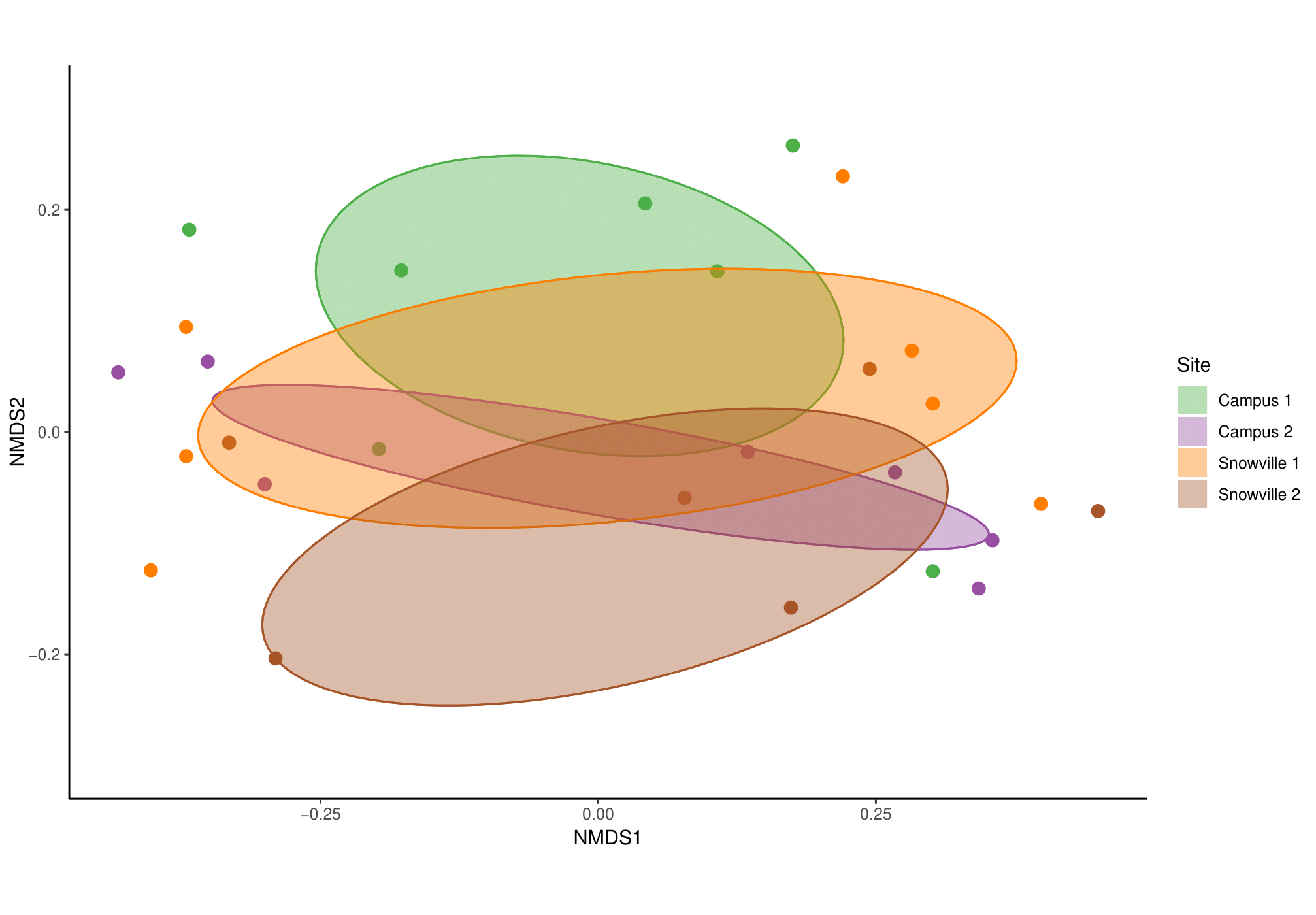

**Figure S3:** Non-metric multidimensional scaling representing insect community composition at four collection sites from both malaise and fermentation traps combined. Ellipsoids represent 95% confidence of the mean for collection site. Points displayed represent community composition for each sample.
